# Spatial organization of adenylyl cyclase and its impact on dopamine signaling in neurons

**DOI:** 10.1101/2023.12.06.570478

**Authors:** Léa Ripoll, Mark von Zastrow

**Affiliations:** Department of Psychiatry and Behavioral Sciences, University of California, San Francisco, San Francisco, CA, USA; Department of Cellular and Molecular Pharmacology, University of California, San Francisco, San Francisco, CA, USA; Quantitative Biology Institute, University of California, San Francisco, San Francisco, CA, USA

## Abstract

The cAMP cascade is widely recognized to transduce its physiological effects locally through spatially limited cAMP gradients. However, little is known about how the adenylyl cyclase enzymes, which initiate cAMP gradients, are localized. Here we answer this question in physiologically relevant striatal neurons and delineate how AC localization impacts downstream signaling functions. We show that the major striatal AC isoforms are differentially sorted between ciliary and extraciliary domains of the plasma membrane, and that AC9 is uniquely targeted to endosomes. We identify key sorting determinants in the N-terminal cytoplasmic domain responsible for isoform-specific localization. We also show that AC9-containing endosomes accumulate activated dopamine receptors and form an elaborately intertwined network with juxtanuclear PKA stores bound to Golgi membranes. Finally, we show that endosomal localization is critical for AC9 to selectively elevate PKA activity in the nucleus relative to the cytoplasm. These results reveal a precise spatial landscape of the cAMP cascade in neurons and a key role of AC localization in directing downstream signal transduction to the nucleus.

## Introduction

Cyclic AMP (cAMP) is a pivotal second messenger that transduces extracellular chemical cues recognized by G protein-coupled receptors (GPCRs) on the cell surface into downstream physiological responses inside the cell. Despites its diffusible nature, cAMP is widely recognized to transduce many of its physiological effects locally via spatially limited cAMP gradients^1^. Local cAMP signaling is not restricted to the plasma membrane, as GPCRs can be activated from different subcellular locations and produce distinct downstream cellular responses^2–12^. To date, the generation of cAMP compartments has been attributed predominantly to phosphodiesterases (PDEs) that restrict cAMP diffusion and to A-kinase anchoring proteins (AKAPs) that sequester cAMP-dependent protein kinase (PKA) and other signaling enzymes within nanodomains^13–16^. However, a key prerequisite for local signaling to occur is the local production of cAMP, but relatively little is known about how the adenylyl cyclase (AC) enzymes that produce cAMP are localized within cells.

Nine homologous AC isoforms (AC1-AC9) mediate GPCR-regulated cAMP production in mammals. All are integral membrane proteins whose enzymatic activity is stimulated by the heterotrimeric G proteins Gα_s/olf_ but they differ in additional regulation by other biochemical inputs, such as Gα_i/o_ proteins and calcium^17^. Much of what is presently known about AC localization concerns AC partitioning at the plasma membrane between lipid domains (‘rafts’ *vs* ‘non-rafts’)^18,19^ and scaffolding into signaling domains by binding to AKAPs^20,21^. An accumulating body of evidence indicates that AC isoforms are also functionally organized at internal membrane locations^2,4,22–26^, but how such isoform-specific localization at distinct membranes is achieved in physiologically relevant cells remains largely unknown.

Neurons represent a relevant cell type in which local cAMP signaling has been shown to occur and to impact downstream physiology^27–33^. In the striatum, dopamine regulates motor and motivated behaviors through cAMP-mediated control of PKA activity in medium spiny neurons (MSNs). Interestingly, disrupting PKA compartmentalization by knocking-out PKA RIIβ, the most abundant striatal regulatory subunit isoform, specifically blocks PKA-mediated regulation of neuronal gene expression without impairing acute locomotor effects induced by dopamine^34^. This highlights the role of local PKA activation in regulating long-term neural adaptations required for motor learning^35^.

Neurons express more than one AC isoform with distinct regulatory properties^36,37^. In the striatum, AC5 is responsible for the majority of the cAMP production^38^ but AC3 and AC9 are also expressed^37,39,40^. Interestingly, AC5 knock-out in mouse striatum blocks some but not all physiological effects of dopamine^41^, and selectively impairs some but not all forms of striatum-dependent learning^42^. Together, these observations raise the questions of whether these isoforms are selectively distributed within cells, and whether differences in their subcellular localization confer distinct effects on downstream functions.

Here, we delineate a precise spatial landscape of isoform-specific localization in MSNs and provide mechanistic insight to how these differences are determined. Focusing on AC9, which is uniquely targeted to endosomes, we reveal an essential role of AC sorting within the endosomal network in directing downstream control of PKA activity to the nucleus.

## Results

### Isoform-specific targeting of striatal ACs in the plasma membrane and endosomes

To begin investigating the subcellular distribution of the selected striatal AC isoforms, we constructed N-terminally HA-tagged versions of each isoform and visualized their subcellular localization in cultured striatal MSNs. Confocal imaging of fixed cells localized HA-AC3 predominantly on the neuronal cell body in a hair-like microdomain that we identified as the primary cilium by colocalization with the ciliary membrane marker Arl13b (**Fig. 1a**), consistent with previous reports^40,43^. HA-AC5 localized in the ciliary microdomain on the soma but was also present in the extraciliary plasma membrane, on the soma and processes (**Fig. 1b**). HA-AC9, in contrast, was distributed on the neuron surface exclusively outside of the ciliary microdomain and also localized to intracellular puncta (**Fig. 1c**). Ciliary localization of HA-AC3 and HA-AC5 was observed in over 90% of neurons examined but was not detected with HA-AC9 (**Fig. 1d**). Fluorescence intensity analysis verified selective enrichment of both AC3 and AC5, but not AC9, in the ciliary microdomain, with AC3 concentrating more strongly than AC5 (**Fig. 1e**).

**Fig 1.**
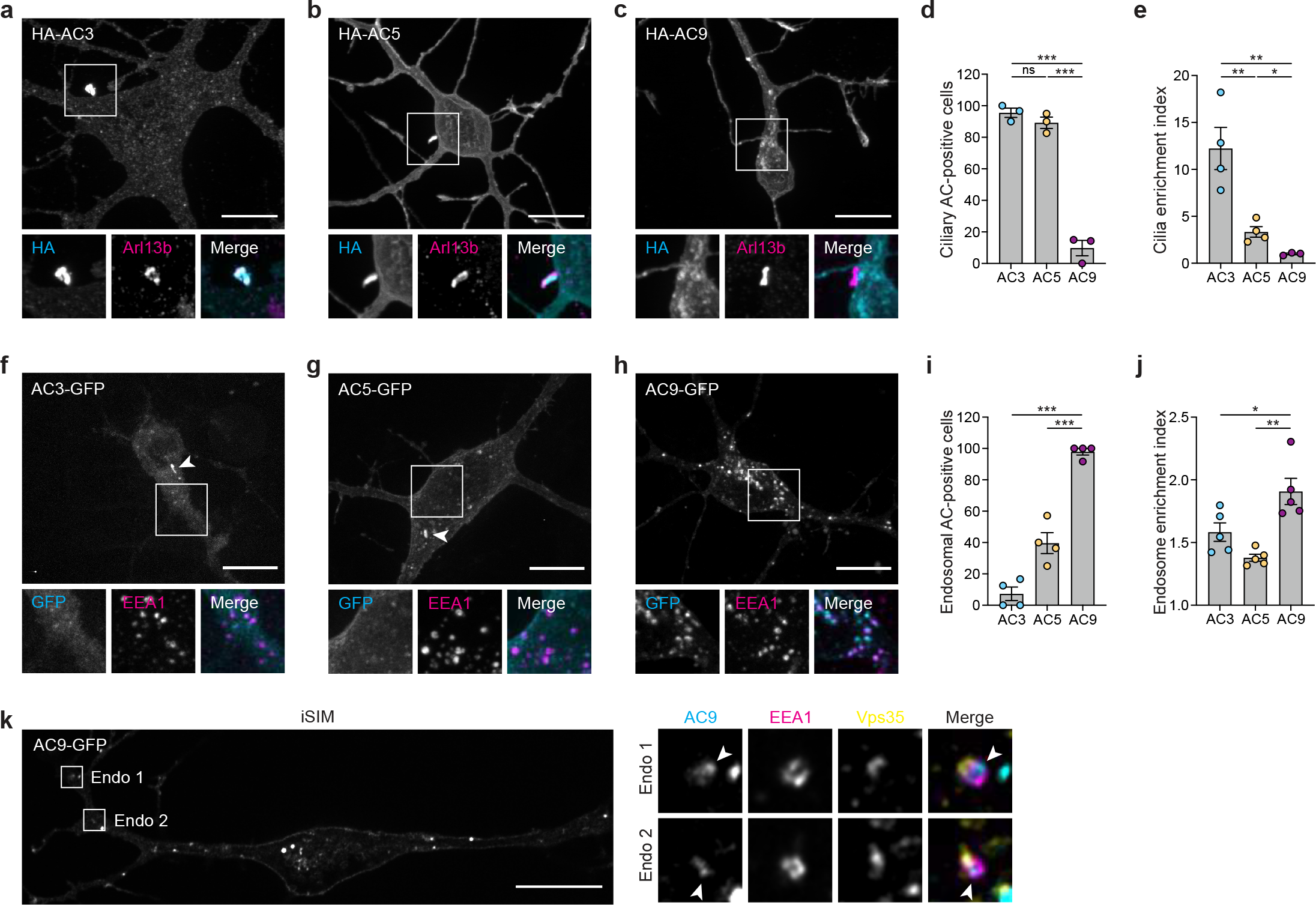
Adenylyl cyclases isoforms localize to primary cilia and endosomes. **a-c**, Maximum intensity Z-projection of confocal microscopy images of MSNs expressing HA-AC3 (**a**), HA-AC5 (**b**) and HA-AC9 (**c**) and stained for cilia marker Arl13b. **d**, Quantification of cilia localization by measuring the fraction of neurons with AC isoforms localized to cilia, using FLAG-D1R as cilia marker (not shown), and expressed as a percentage of total ciliated cells in the transfected cell population. Data are shown as mean ± s.e.m from *n* = 3. **e**, Cilia enrichment index calculated as a ratio of HA-AC fluorescence intensity in cilia (determined by Arl13b and FLAG-D1R) divided by total cell fluorescence. Data are shown as mean ± s.e.m from *n* = 4 (for AC3 and AC5) and *n* = 3 (for AC9). **f-h**, Representative confocal microscopy images of MSNs expressing AC3-GFP (**f**), AC5-GFP (**g**) and AC9-GFP (**h**) and stained for endosomal marker EEA1. Arrowheads indicate cilia. **i**, Quantification of endosome localization by measuring the fraction of neurons with 10 or more AC-GFP internal puncta, and expressed as a percentage of total transfected cells. Data are shown as mean ± s.e.m from *n* = 4. **j**, Endosome enrichment index calculated as a ratio of AC-GFP fluorescence intensity at EEA1 positive endosomes divided by total cell fluorescence. Data are shown as mean ± s.e.m from *n* = 5. **k**, Representative iSIM images of MSN expressing AC9-GFP and stained for EEA1 and Vps35. Arrowheads indicate AC9-positive subdomains at the endosomal membrane. Scale bars are 10 μm. \*\*\**P*<0.001, \*\**P*<0.01, \**P*<0.05, n.s. not significant by unpaired two-tailed Student’s *t*-test.

We confirmed these isoform-specific subcellular distribution patterns using AC constructs labeled in the C-terminus rather than N-terminus with a monomeric GFP variant^44^ (muGFP) rather than an epitope tag (**Fig. 1f-h**, arrowheads in each image indicate the cilium). Interestingly, the vast majority of AC9-containing intracellular puncta overlapped with EEA1 (**Fig. 1h**, lower panels) and Vps35 (**Extended Data Fig. 1a**), identifying them as endosomes. Robust intracellular accumulation of AC9 was observed in almost all neurons examined (**Fig. 1i**) and we verified the selective endosomal enrichment of AC9 by fluorescence intensity measurement using EEA1 (**Fig. 1j**) or Vps35 (**Extended Data Fig. 1b**) to define an endosome mask. Using instant Structured Illumination Microscopy (iSIM) to obtain higher spatial resolution, we found AC9-GFP to be concentrated on subdomains of the endosomal limiting membrane (**Fig. 1k**, arrowheads) and, in some endosomes, it appeared to be in the endosome lumen (**Extended Data Fig. 1c**).

Altogether, these data reveal distinct subcellular distribution of AC isoforms in neurons. AC3 is essentially restricted to the surface of the soma through concentration in the ciliary microdomain. AC5 localizes on the soma both in cilia and on the extraciliary surface, extending into dendritic and axonal processes. AC9 localizes on the neuronal plasma membrane exclusively outside of cilia and is selectively concentrated in endosomes where it appears to be organized in membrane microdomains.

### D1Rs colocalize with ACs at each isoform-selective membrane location

GPCR-mediated stimulation of AC activity is a membrane-delimited process thought to require the presence of activated receptors and ACs in the same bilayer^45^. MSNs exhibit high expression of the dopamine 1 receptor (D1R), a Gα_s/olf_-coupled GPCR whose activation can also occur at endosomes, Golgi, and primary cilia, resulting in distinct downstream outcomes^4,46,47^. Accordingly, we next investigated the subcellular localization of each isoform relative to D1R. Confirming previous results^48,49^, FLAG-tagged D1R localized at the plasma membrane but also in the ciliary microdomain together with HA-AC3 (**Fig. 2a**) and HA-AC5 (**Fig. 2b**).

**Fig 2.**
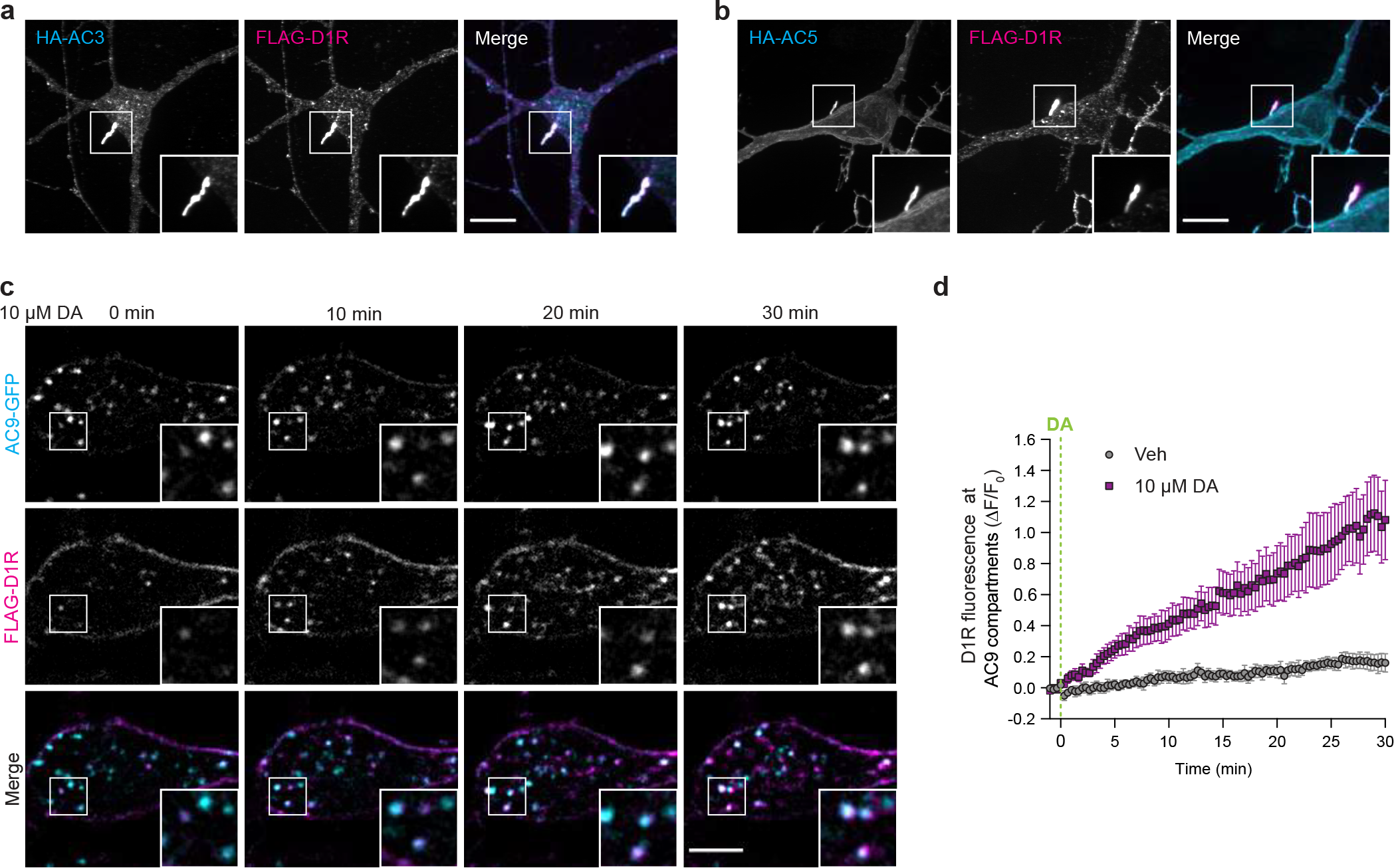
Dopamine 1 receptor localizes to adenylyl cyclase isoforms positive compartments. **a,b**, Maximum intensity Z-projection of confocal microscopy images of MSNs expressing FLAG-D1R and HA-AC3 (**a**) or HA-AC5 (**b**). Scale bar = 10 μm. **c**, Representative live cell spinning disk confocal images of MSNs transfected with AC9-GFP and FLAG-D1R and treated with 10 μM dopamine (DA) at 0 min. Surface FLAG-D1R was labeled with Alexa Fluor 555-coupled anti-FLAG antibody for 15 min before imaging. Scale bar = 5 μm. **d**, Quantification of surface labeled FLAG-D1R accumulation at segmented AC9-GFP-positive endosomes after vehicle (Veh) or 10 μM DA addition. Data are shown as mean ± s.e.m from *n* = 5.

In contrast to D1R targeting to the primary cilium, D1R localization in endosomes is agonist-dependent and requires agonist-induced endocytosis and subsequent delivery to endosomes^50,51^. We verified this and used live cell imaging to ask if D1Rs accumulate in the same or different endosomes as AC9. Surface-labeled FLAG-D1Rs redistributed from the plasma membrane to internal membrane puncta within minutes after addition of 10 μM dopamine to the culture medium. Further, the internalized D1Rs were found to accumulate in the same puncta containing AC9 (**Fig. 2c**), defined as endosomes by colocalization with EEA1 (**Extended Data Fig. 2a,b**). Fluorescence intensity analysis revealed that D1Rs begin to accumulate in AC9-containing endosomes within ∼1 min after dopamine application and then progressively accumulate over a prolonged time course (**Fig. 2d**). The localization of AC9 to endosomes, in contrast, remained stable (**Extended Data Fig. 2c,d**).

Altogether, these results indicate that D1Rs can populate each of the subcellular membrane domains in which striatal AC isoforms distribute: D1R localizes in the plasma membrane, both to the ciliary microdomain containing AC3 and AC5 and to the extraciliary domain containing AC5 and AC9. D1R can also robustly accumulate in endosomes containing AC9, but this process is ligand-dependent.

### The N-terminal cytoplasmic domain harbors isoform-selective targeting determinants

Having established distinct subcellular localization patterns among the AC isoforms examined, we next investigated the structural basis for the isoform-selective targeting. Transmembrane ACs have a shared membrane topology and share extensive sequence homology, but the N-terminal cytoplasmic domain of ACs is highly divergent^17^. Accordingly, we focused on this domain as a potential determinant of isoform-selective AC targeting.

To examine AC targeting to the primary cilium, we assessed the effect of replacing the N-terminus of AC5, which selectively concentrates in the primary cilium but is also detectable in the plasma membrane outside of the cilium, with the corresponding sequence derived from the non-ciliary isoform AC9 (**Fig. 3a**). We found that this chimeric mutant protein (AC5-AC9-Nter), while still observed in the plasma membrane, failed to localize on the primary cilium, labeled with FLAG-D1R (**Fig. 3a,b**). Deleting the N-terminus from AC5 (AC5-ΔNter) produced a similar phenotype (**Fig. 3c,d**). Together, these results indicate that the N-terminal cytoplasmic domain of AC5 is necessary for targeting this isoform to the cilium. Conversely, replacing the N-terminus of AC9 with the corresponding sequence derived from AC5 produced a chimeric mutant construct (AC9-AC5-Nter) that accumulated in the primary cilium (**Fig. 3e,f**). This indicates that the N-terminal cytoplasmic domain of AC5 is also sufficient to drive AC targeting to the cilium.

**Fig 3.**
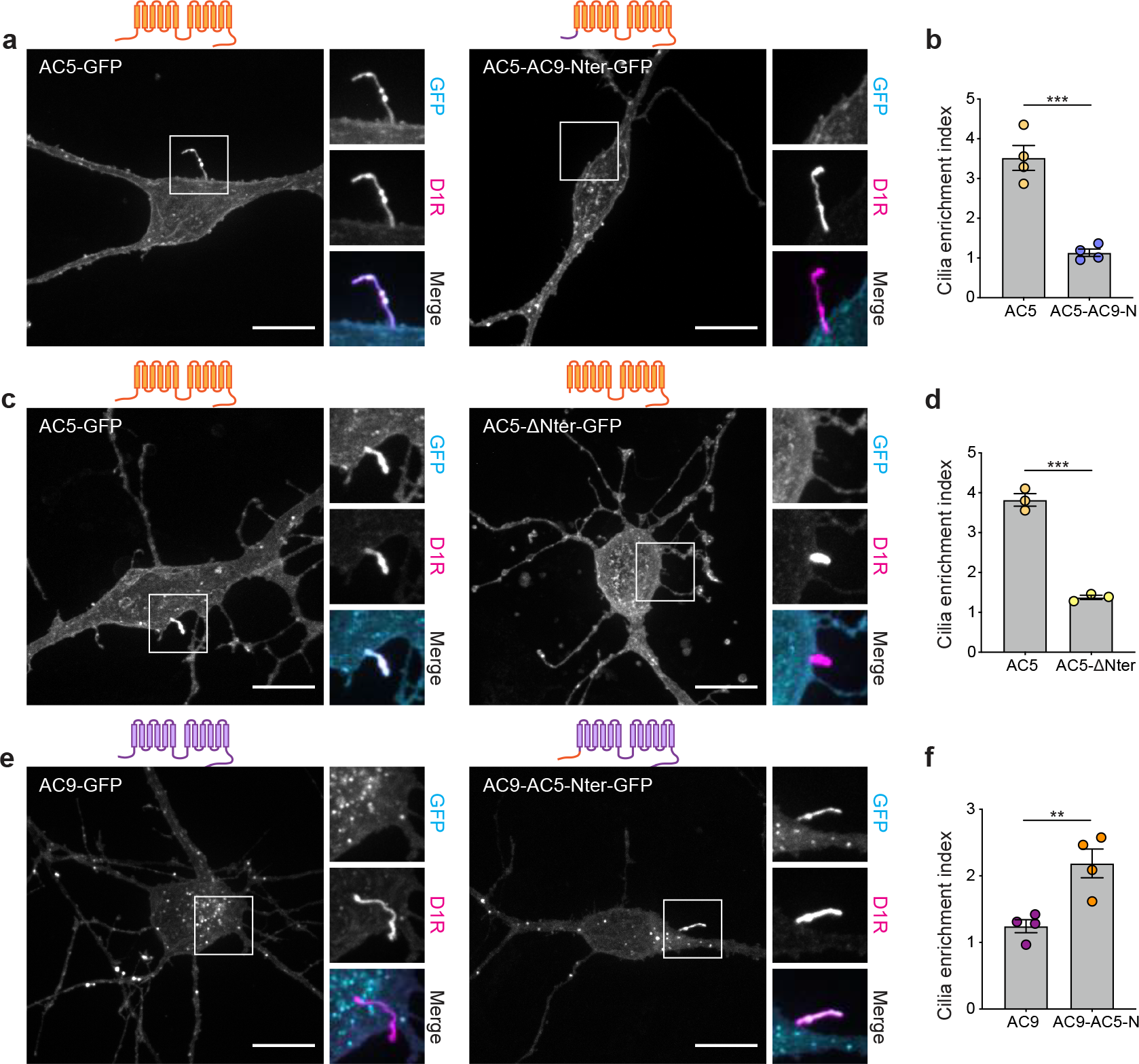
The N-terminus of AC5 is necessary and sufficient for cilia targeting. **a,c,e**, Maximum intensity Z-projection of confocal images microscopy images of MSNs surface-labeled for FLAG-D1R and expressing AC5-GFP or AC5-AC9-Nter-GFP (**a**), AC5-GFP or AC5-ΔNter-GFP (**c**), AC9-GFP or AC9-AC5-Nter-GFP (**e**). Show above each panel is the schematic representation of each construct showing corresponding mutations and topology. **b,d,f**, Cilia enrichment index of cells co-expressing FLAG-D1R and AC5-GFP or AC5-AC9-Nter-GFP (**b**, *n* = 4), AC5-GFP or AC5-ΔNter-GFP (**d**, *n* = 3), AC9-GFP or AC9-AC5-Nter-GFP (**f**, *n* = 4). Data are shown as mean ± s.e.m. Scale bars are 10 μm. \*\*\**P*<0.001, \*\**P*<0.01 by unpaired two-tailed Student’s *t*-test.

We noticed above that the chimeric mutant AC5-AC9-Nter (**Fig. 4a**) localized to intracellular puncta (**Fig. 3a**). To further investigate the structural determinants responsible for endosome targeting, we verified AC5-AC9-Nter accumulation in endosomes by colocalization with EEA1 (**Fig. 4b**). Using fluorescence intensity analysis, we determined that fusing the AC9 N-terminus to AC5 promotes endosomal concentration of the chimeric mutant to the same degree as observed for AC9 (**Fig. 4c**). These results indicate that the N-terminal cytoplasmic domain of AC9 is sufficient to drive isoform-selective AC targeting to endosomes.

**Fig 4.**
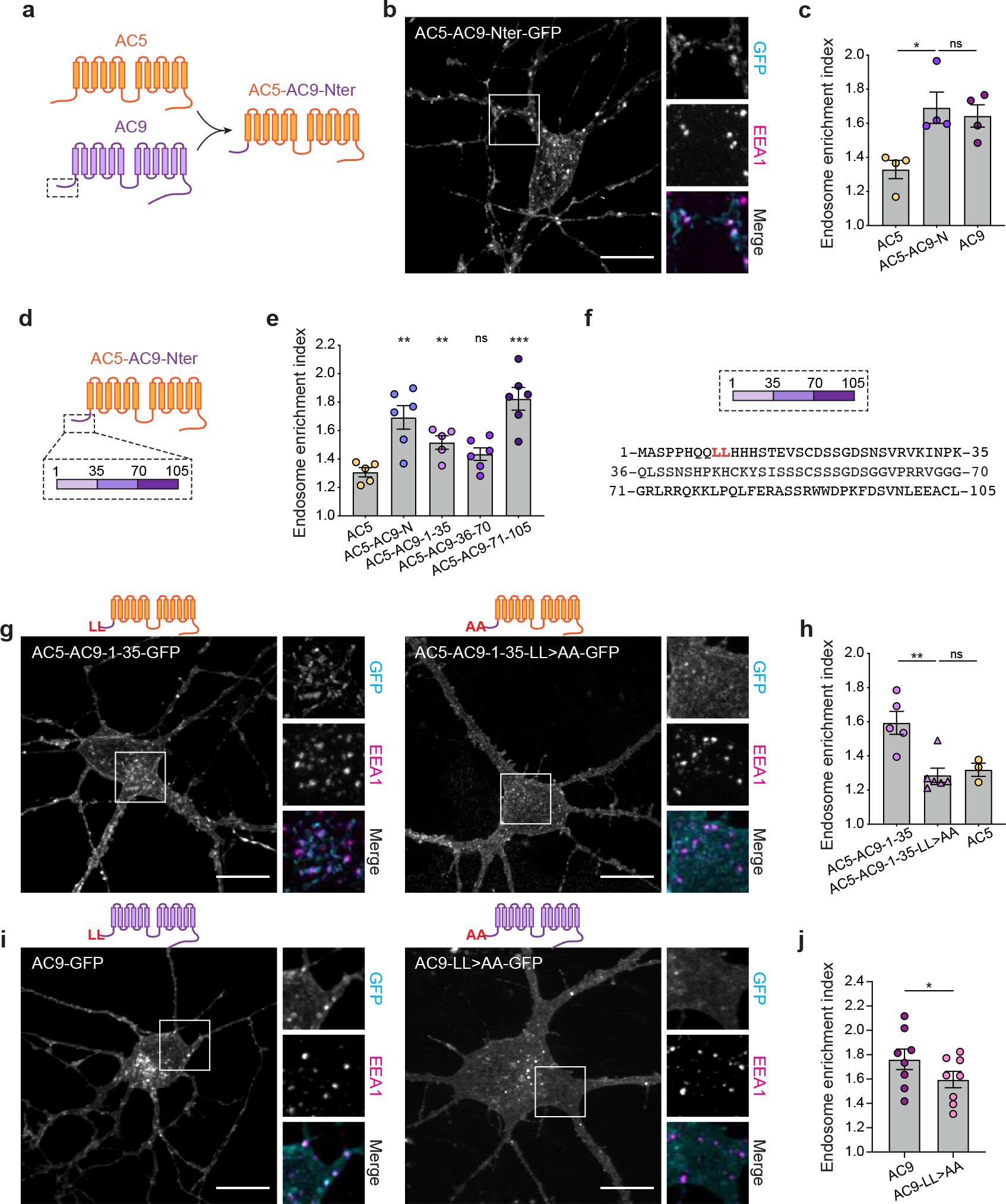
A dileucine motif in the N-terminus of AC9 is required for endosome localization. **a**, Schematic representation of the chimeric mutant AC5-AC9-Nter. AC5-derived sequence is depicted in orange and AC9-derived sequence in purple. **b**, Maximum intensity Z-projection of confocal images microscopy images of MSNs expressing AC5-AC9-Nter-GFP and stained for EEA1. **c**, Endosome enrichment index of cells expressing AC5-GFP, AC5-AC9-Nter-GFP or AC9-GFP. Data are shown as mean ± s.e.m from *n* = 4. **d**, Schematic of chimeric mutants of AC5 (in orange) containing portions of AC9 N-terminus (in purple). **e**, Endosome enrichment of cells expressing AC5 chimeric mutants. Data are shown as mean ± s.e.m from *n* = 5-6. **f**, Schematic of the three portions of AC9 N-terminus and their respective sequence. **g**, Representative confocal images of MSNs expressing AC5-AC9-1-35-GFP or AC5-AC9-1-35-LL>AA-GFP and stained for EEA1. **h**, Endosome enrichment of cells in **g**. Data are shown as mean ± s.e.m from *n* = 3-6. **i**, Representative confocal images of MSNs expressing AC9-GFP or AC9-LL>AA-GFP and stained for EEA1. **j**, Endosome enrichment of cells in **i**. Data are shown as mean ± s.e.m from *n* = 8. Scale bars are 10 μm. \*\**P*<0.01, \**P*<0.05, n.s. not significant by unpaired two-tailed Student’s *t*-test.

### A dileucine motif in the AC9 N-terminus drives robust concentration in endosomes

We sought to map the endosomal targeting determinant in the AC9 N-terminal domain in more detail by dividing this sequence into three portions and examining the ability of each to drive chimeric AC5 targeting to endosomes (**Fig. 4d**). The middle portion of the AC9 N-terminal tail (amino acids 36-70) did not promote AC5 localization to endosomes, but both the proximal (amino acids 1-35) and distal (amino acids 71-105) portions conferred detectable endosomal enrichment on AC5 chimeras (**Fig. 4e**). We noticed that the proximal portion contains a dileucine motif (**Fig. 4f**, highlighted in red), which is widely known to target membrane proteins to endosomes by promoting their endocytosis via clathrin-coated pits^52^. Thus, we asked if the dileucine motif present in the AC9 N-terminus also acts as an endocytic determinant for this isoform. Mutating the dileucine residues to alanines within the AC5 chimera containing AC9 N-terminus proximal portion (AC5-AC9-1-35-LL>AA) led to a complete loss of endosomal localization (**Fig. 4g,h**). Furthermore, mutating this dileucine motif in full-length AC9 (AC9-LL>AA) significantly reduced AC9 enrichment in endosomes (**Fig. 4i,j**), causing a shift in the distribution of AC9 from endosomes to the plasma membrane. However, this mutation did not fully block visible AC9 targeting to endosomes, consistent with the distal portion of AC9 N-terminus also having endocytic activity. Together, these results indicate that the dileucine motif present in the proximal segment of AC9 N-terminus acts as part of a bipartite endosomal targeting and is required for robust endosomal concentration of AC9.

### Endosomal localization of AC9 impacts the timing of dopamine-induced PKA activity

We next investigated the significance of AC subcellular localization by assessing the impact of each isoform on the cAMP/PKA signaling cascade. AC3 and AC5 overexpression significantly elevated levels of both global cAMP concentration and PKA activity elicited by endogenous dopamine receptor activation (**Fig. 5a,b**). Surprisingly, AC9 overexpression did not detectably increase global cAMP but it significantly augmented the elevation of PKA activity elicited by endogenous dopamine receptors (**Fig. 5c**).

**Fig 5.**
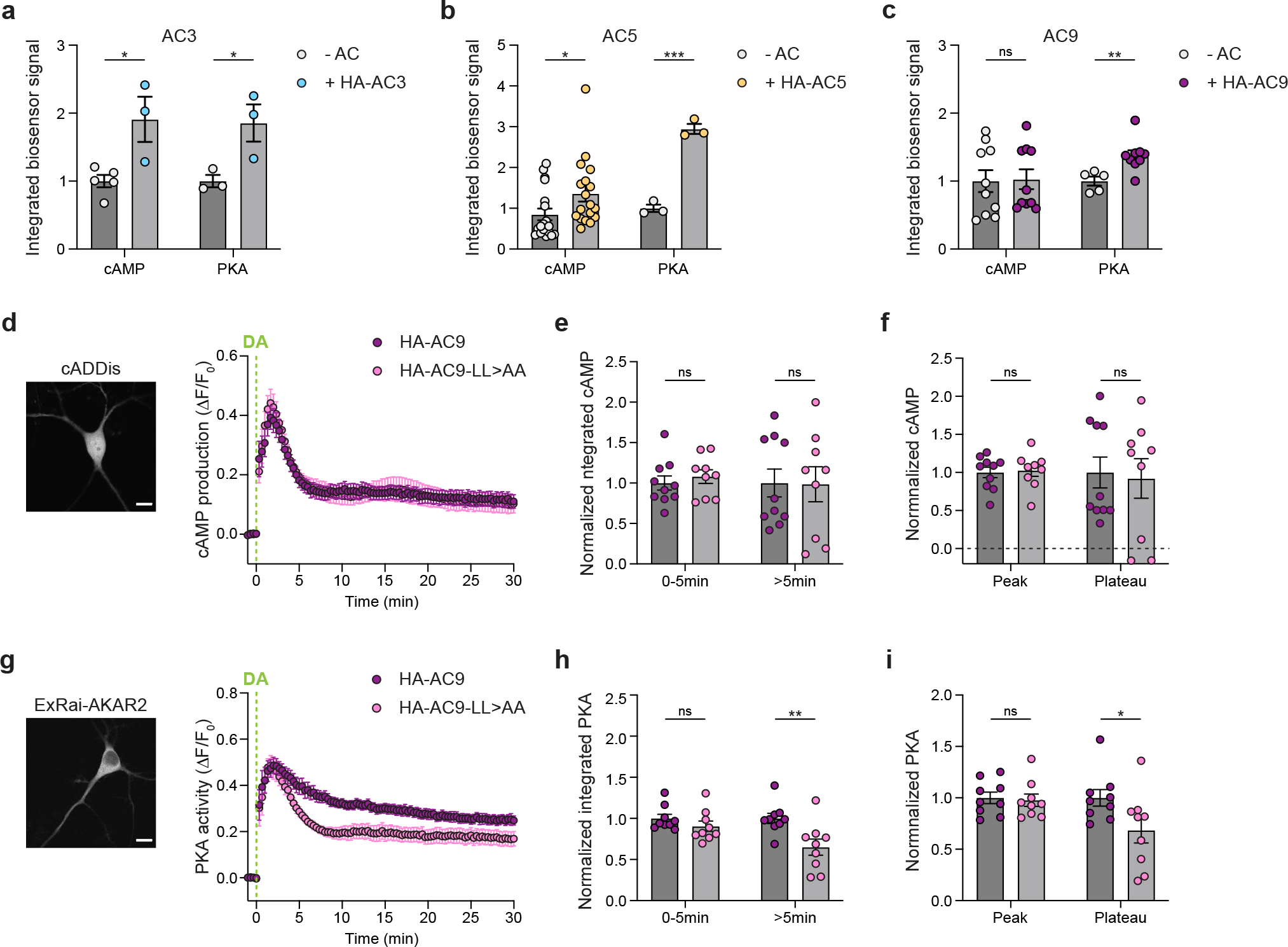
Endosomal AC9 modulates PKA activity but not overall cAMP. **a-c**, Forskolin (Fsk) and IBMX-normalized cAMP and PKA response in cells expressing the cADDis (cAMP) or ExRai-AKAR2 (PKA) sensors alone or with HA-AC3 (**a**), HA-AC5 (**b**) or HA-AC9 (**c**) and treated with 10 μM dopamine (DA). Data are shown as mean ± s.e.m. (**a**) *n* = 5 and 3 (cAMP, -AC and HA-AC3, respectively) and *n* = 3 (PKA); (**b**) *n* = 18 (cAMP) and *n* = 3 (PKA); (**c**) *n* = 10 (cAMP) and *n* = 5 and 9 (PKA, -AC and HA-AC9 respectively). **d**, On the left, representative image of a cell expressing the cAMP biosensor cADDis. On the right, kinetics of cAMP production over time in MSNs coexpressing cADDis and HA-AC9 (purple, *n* = 10) or HA-AC9-LL>AA (pink, *n* = 9) and treated with 10 μM DA. The ΔF/F0 was measured every 20 sec and normalized to the Fsk and IBMX response (added after 30 min of DA treatment). **e**, Integrated cAMP signals of the phases 0-5 min and >5 min (5-30 min) after DA addition were calculated as the area under the curve and normalized to the average HA-AC9 value. **f**, Peak and plateau values were calculated as the maximum ΔF/F0 (peak) and the average of 20-30 min values (plateau) and were normalized to the HA-AC9 value. **g**, On the left, representative image of a cell expressing the PKA biosensor ExRai-AKAR2. On the right, kinetics of PKA activity over time in MSNs coexpressing ExRai-AKAR2 and HA-AC9 (purple, *n* = 9) or HA-AC9-LL>AA (pink, *n* = 9) and treated with 10 μM DA. The ΔF/F0 was measured every 20 sec and normalized to the Fsk and IBMX response. **h**, Integrated PKA signals of the phases 0-5 min and >5 min (5-30 min) after DA addition were calculated as the area under the curve and normalized to the average HA-AC9 value. **i**, Peak and plateau values were calculated as the maximum ΔF/F0 (peak) and the average of 20-30 min values (plateau) and were normalized to the HA-AC9 value. For all panels, data represent biological replicates and are shown as mean ± s.e.m. Scale bars are 10 μm. \*\*\**P*<0.001, \*\**P*<0.01, \**P*<0.05, n.s. not significant by unpaired two-tailed Student’s *t*-test.

To ask if this selective effect of AC9 on PKA relative to cAMP is related to AC9 concentration in endosomes, we used the dileucine mutation (AC9-LL>AA) to selectively reduce endosomal localization (**Fig. 4i,j**). Notably, the expression of AC9-LL>AA did not impact the overall endogenous dopamine-induced cAMP production when compared to AC9 expression. In both conditions, a rapid cAMP elevation occurred within 2 minutes and decayed over several minutes to a plateau above baseline in the continuous presence of agonist (**Fig. 5d**). Both the rapid initial elevation and the plateau phase remained unaltered by AC9-LL>AA expression (**Fig. 5e,f**). Subsequently, we evaluated the impact of AC9 mis-localization on endogenous dopamine-elicited elevation of global PKA activity using the ExRai-AKAR2 fluorescent biosensor^53^. Surprisingly, while the initial peak of PKA activity remained unchanged between the two conditions, AC9-LL>AA was significantly impaired relative to wild type AC9 in its ability to support the plateau phase of PKA activity elevation (**Fig. 5g-i**).

These observations suggest that the endosomal localization of AC9 impacts the timing of the PKA response, and specifically the ability to maintain the later plateau phase of activity elevation. However, the effect is not attributable to any detectable increase in global cAMP levels. This led us to investigate in more detail the spatial relationship in neurons between AC9-containing endosomes and PKA.

### AC9-containing endosomes dynamically contact juxtanuclear neuronal PKA stores

PKA is a heterotetramer consisting of two catalytic subunits and two regulatory subunits, with PKA regulatory subunits IIβ (PKA RIIβ) being the main isoforms in the striatum^34,54–56^. Superresolution iSIM microscopy analysis revealed close proximity and intertwining of AC9-positive endosomes with both PKA catalytic (PKA cat) (**Fig. 6a,b** and **Supplementary Video 1**) and PKA RIIβ (**Fig. 6c,d** and **Supplementary Video 2**) compartments. These tubulovesicular PKA cat and RIIβ compartments were localized in the perinuclear region in a distribution overlapping that of Golgi membranes (**Extended Data Fig. 3a,b**), as previously observed^9,57–62^. Furthermore, we verified extensive colocalization between PKA cat and RIIβ in the Golgi region (**Extended Data Fig. 3c**), consistent with complex formation there. Live cell microscopy of neurons revealed dynamic close interactions between AC9-positive endosomes and PKA cat puncta, following dopamine addition, with contact times ranging from several minutes up to ∼30 minutes (**Fig. 6e, Supplementary Video 3** and **Extended Data Fig. 4**). Close contact between AC9-positive endosomes and perinuclear PKA cat stores was suggested by coordinated movement of the two adjacent proteins in sequential image series (**Fig. 6f**).

**Fig 6.**
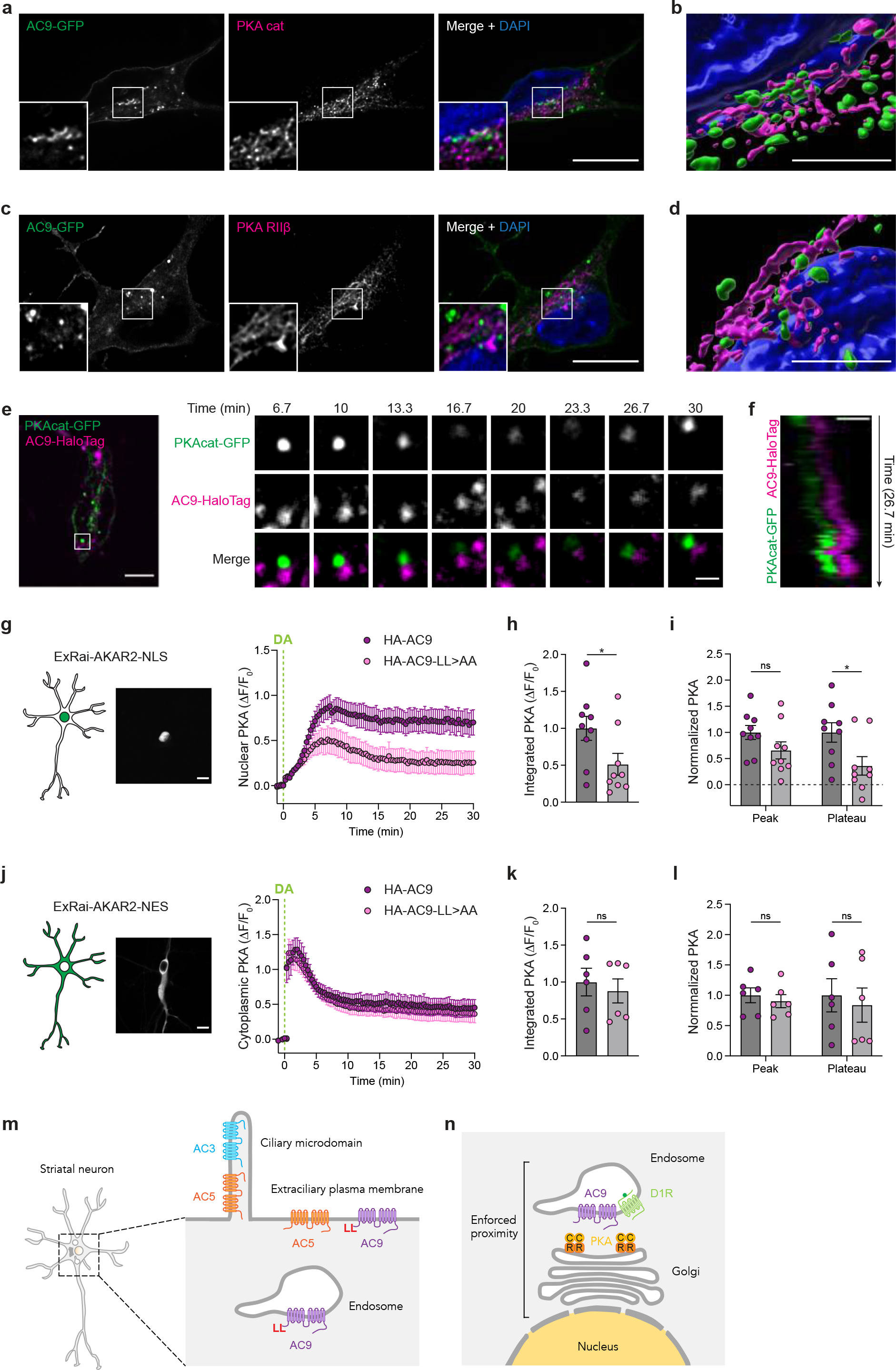
Nuclear PKA activity is dependent on AC9 endosomal concentration. **a,c**, Representative iSIM images of MSN expressing AC9-GFP and stained for endogenous PKA cat (**a**) orPKA RII (**c**). Nuclei were stained with DAPI. Scale bar = 10 μm. **b,d**, 3D rendering of cells in **a** (**b**) and **c** (**d**). Scale bar = 5 μm. **e**, Spinning-disk confocal images from a time series of neurons coexpressing PKAcat-GFP and AC9-HaloTag and treated with 10 μM dopamine (DA) at t =0 min. Scale bar = 5 μm. **f**, Kymograph of the cell in **e**. PKA cat puncta and AC9-containing endosomes move together in the soma of MSNs. Scale bar = 1 μm. **g**, On the left, schematic representation and spinning-disk confocal representative image showing ExRai-AKAR2-NLS localization to the nucleus (Scale bar = 10 μm). On the right, kinetics of nuclear PKA activity over time in MSNs coexpressing ExRai-AKAR2-NLS and HA-AC9 (purple, *n* = 9) or HA-AC9-LL>AA (pink, *n* = 9) and treated with 10 μM dopamine. The ΔF/F0 was measured every 20 sec. **h**, Integrated nuclear PKA signal was calculated as the area under the curve and normalized to the average HA-AC9 value. **i**, Peak and plateau values were calculated as the maximum Δ F/F0 (peak) and the average of 20-30 min values (plateau) and normalized to the HA-AC9 value. **j**, On the left, schematic representation and spinning-disk confocal representative image showing ExRai-AKAR2-NES localization to the cytoplasm and excluded from the nucleus (Scale bar = 10 μm). On the right, kinetics of cytoplasmic PKA activity over time in MSNs coexpressing ExRai-AKAR2-NES and HA-AC9 (purple, *n* = 6) or HA-AC9-LL>AA (pink, *n* = 6) and treated with 10 μM dopamine. The ΔF/F0 was measured every 20 sec. **k**, Integrated cytoplasmic PKA signal was calculated as the area under the curve and normalized to the average HA-AC9 value. **l**, Peak and plateau values were calculated as the maximum Δ F/F0 (peak) and the average of 20-30 min values (plateau) and normalized to the HA-AC9 value. **m**, Diagram of proposed model illustrating the distinct subcellular distribution of AC isoforms in striatal neurons. AC3 is restricted to the primary cilium; AC5 localizes at the plasma membrane, both in cilia and extraciliary surfaces; AC9 localizes at the plasma membrane outside of cilia, and in endosomes. Endosomal localization requires a dileucine motif in AC9 N-terminus. **n**, Endosomes containing AC9 and D1R are in close proximity with Golgi-associated PKA compartments in the perinuclear region. This enforced proximity ‘in trans’ selectively drives PKA activity in the nucleus (yellow). For all panels, data represent biological replicates and are shown as mean ± s.e.m. \**P*<0.05, n.s. not significant by unpaired two-tailed Student’s *t*-test.

### Concentration in endosomes enables AC9 to specifically control nuclear PKA activity

A prevailing current view of how cAMP elevates nuclear PKA activity is by inducing dissociation of catalytic subunits from regulatory subunits, disinhibiting the kinase activity of catalytic subunits and enabling a fraction of them to diffuse into the nucleus^63–66^. Based on the close apposition between AC9 endosomes and PKA compartments in the perinuclear region, we wondered whether the ability of AC9 endosomal localization to increase PKA activity could reflect an effect of PKA activity elevation in the nucleus. To test this, we generated a nucleus localized-ExRai-AKAR2 (ExRai-AKAR2-NLS) and co-expressed it with AC9 or AC9-LL>AA in neurons (**Fig. 6g**). Dopamine addition elicited a gradual increase in nuclear PKA activity that peaked between 5 to 10 minutes, followed by a plateau phase (**Fig. 6g**). Both the total nuclear PKA activity (**Fig. 6h**) and the plateau phase were strongly reduced upon AC-LL>AA expression (**Fig. 6i**). As an alternative approach, we measured nuclear PKA activity using the non-targeted ExRai-AKAR2 (**Fig. 5g**) by drawing a ROI within the nucleus and measuring fluorescence over time (**Extended Data Fig. 5a**). AC9-LL>AA expression similarly resulted in a significant reduction in nuclear PKA activity (**Extended Data Fig. 5b,c**), affecting both the peak and plateau phases (**Extended Data Fig. 5d**). These results suggest a critical role for endosomal concentration of AC9 in promoting the ability of endogenous dopamine receptor activation to elevate PKA activity in the nucleus.

We next carried out the converse analysis, constructing an ExRai-AKAR2 variant that is diffusely localized in the cytoplasm and excluded from the nucleus (ExRai-AKAR2-NES, **Fig. 6j**). Remarkably, and in contrast to the clear difference observed for PKA activity elevation in the nucleus, reducing AC9 concentration in endosomes (AC9-LL>AA) produced no discernible effect on the dopamine-elicited elevation of PKA activity in the cytoplasm (**Fig. 6k,l**). Altogether, our data collectively reveal that AC9 endosomal localization specifically and uniquely modulates PKA activity by promoting PKA activity elevation in the nucleus, without detectably affecting cytoplasmic activity.

## Discussion

Local cAMP signaling from any subcellular location requires cAMP to be locally produced, but a key knowledge gap in our present understanding is how ACs are localized. We addressed this by focusing on striatal MSNs, a native cell type in which the cAMP/PKA cascade mediates most of the physiological actions of dopamine^67–69^. We describe a precise spatial organization of striatum-expressed AC isoforms in these neurons, delineate a discrete cellular mechanism for targeting AC9 to endosomes, and reveal an elaborate membrane network in the somatic cytoplasm that enforces proximity between AC9 and PKA in the juxtanuclear cytoplasm. We also show that efficient sorting of AC9 into this network provides a selective way for AC9 to regulate PKA activity in the nucleus.

AC3 has been previously documented to localize to the primary cilium, a limited microdomain of the somatic plasma membrane^40,43^. Our results verify this and reveal additional diversity among isoforms in surface distribution. We show that AC5 also concentrates in the ciliary microdomain but, unlike AC3, it is enriched to a reduced degree and is present also on the extraciliary surface on the soma and dendrites. We then identify the divergent AC N-terminal cytoplasmic domain as a key structural determinant of isoform-selective differences in AC targeting to the primary cilium. AC9 differs from both AC3 and AC5 in being excluded from the ciliary microdomain and specifically concentrated in endosomes. We previously showed AC9 localization to endosomes in HEK293 cells but its trafficking to endosomes relies on ligand-induced activation of the β2-adrenergic receptor and is sensitive to additional environmental cues that remain poorly defined^26^. Here we demonstrate a robust and consistent endosomal localization in relevant cell types that is not dependent on receptor activation. We also define a dileucine motif within the proximal region of AC9 N-terminus that is necessary for its robust endosomal concentration of AC9. In addition, we show that the distal region of AC9 N-terminus also contributes to endosomal enrichment, but the precise motif responsible for this targeting remains to be determined. We note that each of these differentially localized striatal AC isoforms represent distinct AC subclasses, as defined by differential sensitivity to or regulation by G_i/o_ and calcium. AC3 is weakly activated by Ca^2+^/calmodulin^70^, AC5 is inhibited by Gα_i_ and Ca^2+ 71^ and AC9 is insensitive to both^72^. Accordingly, our results delineate a precise spatial landscape of the AC system that places AC isoforms differing in biochemical regulation at distinct subcellular membrane locations (**Fig 6m**).

Here we establish a functional significance of AC sorting in the spatial landscape by focusing on AC9’s unique and specific concentration in endosomes. Previous studies have shown that local cAMP production from internal membranes can confer advantages over the plasma membrane in facilitating signaling in the nucleus^7,9,12,59,62,73^. Our results are fully consistent with this concept and advance the present understanding in several ways. Our study establishes AC9 as a pivotal factor responsible for local cAMP signaling from endosomes, while also uncovering the mechanisms governing AC9 targeting to endosomes. We show that AC9-containing endosomes are closely opposed to juxtanuclear PKA stores by being intimately intercalated with Golgi membranes on which type II PKA is bound. We further show that concentration in this intercalated membrane network enables AC9 to selectively elevate PKA activity in the nucleus relative to the cytoplasm.

Our study emphasizes the intertwined nature of AC-containing endosomes and Golgi-associated PKA compartments, characterized by extensive regions of close association. Local signaling by a diffusible mediator ultimately depends on the principle of enforced proximity^74^. Previous studies have established this principle in cAMP signaling, based on the lateral partitioning of AC and its molecular scaffolding with PKA^20^. This represents an ‘in cis’ form of enforced proximity, based on proximity occurring on the same membrane surface. We propose that the present results reveal a discrete cellular mechanism for achieving enforced proximity ‘in trans’, or between distinct membranes (i.e., endosome and Golgi membranes). The utility of membrane-membrane apposition or contact in achieving enforced proximity in cellular signal transduction is increasingly recognized, such as in signaling by calcium, lipids and reactive oxygen species^75^. To our knowledge, the present results provide the first evidence for extending this general principle to cellular signaling by the cAMP cascade (**Fig 6n**).

Finally, our study reveals the unique ability of AC9 to specifically promote PKA elevation in the nucleus relative to the cytoplasm. This high degree of specificity holds interesting physiological implications. PKA activity within the nucleus is closely linked to long-term cellular changes and learning processes^64^. Therefore, AC9 ability to selectively modulate this nuclear activity carries the potential to significantly impact neuronal function and behavior. It is worth noting that AC9 is widely expressed in the brain, and particularly abundant in the hippocampus, a region associated with learning and memory^76^. Investigating whether AC9 endosomal localization extends to other neuronal cell types and whether it contributes to long-term plasticity associated with learning and memory would be of considerable interest.

## Methods

### Primary rat striatal neuron culture

Medium spiny neurons were prepared from embryonic day 18 Sprague-Dawley rats. After euthanasia of the pregnant Sprague-Dawley rat (CO_2_ and bilateral thoracotomy), the brains of embryonic day 18 rats of both sexes were extracted from the skull. The striatum, including the caudate-putamen and nucleus accumbens, was identified as described^77^. Striata were dissected in ice cold HBSS calcium/magnesium/phenol red free (Thermo Fisher). Structures were dissociated in 0.05% trypsin/EDTA (UCSF Media Production) for 15 min at 37°C and washed in DMEM (Thermo Fisher) supplemented with 10% fetal bovine serum (UCSF Media Production) and 30 mM HEPES. Cells were then mechanically separated with a flame-polished Pasteur pipette and were plated onto poly-D-lysine coated 35 mm glass bottom dishes (Cellvis) in DMEM supplemented with 10% fetal bovine serum. Medium was exchanged on DIV 4-5 for phenol-free Neurobasal medium (Thermo Fisher) supplemented with GlutaMAX 1x (Thermo Fisher) and Gibco B-27 1x (Thermo Fisher) supplements. Half of the culture medium was exchanged every week with fresh, equilibrated medium. Cytosine arabinosine 2 μM (Millipore Sigma) was added at DIV 8. Transfection using Lipofectamine 2000 (Thermo Fisher) was performed at DIV 8 using 2 μl of Lipofectamine and 1-2 μg DNA in 1ml of media per 35 mm imaging dish, and media was exchanged 4-6 hours later. Cells were maintained in a humidified incubator with 5% CO2 at 37°C and imaged at DIV 11-15. All procedures were performed according to the National Institutes of Health Guide for Care and Use of Laboratory Animals and approved by the UCSF Institutional Animal Care and Use Committee (protocol number AN185688).

### Reagents and antibodies

Information on all chemicals and antibodies used in this study can be found in **Supplementary Table 1** and **Supplementary Table 2**.

### DNA constructs

Information on all plasmids used in this study can be found in **Supplementary Table 3**.

N-terminally HA-tagged ACs were generated by PCR amplifying Homo sapiens AC3 cDNA (purchased from DNASU, HsCD00403688), Homo sapiens AC5-EGFP (previously generated in the von Zastrow lab by G. Peng from AC5 cDNA purchased from DNASU, HsCD00732284) and Homo sapiens AC9-EGFP (previously described in ^59^) with N-terminal HA tag added into primers followed by insertion into NotI/EcoRI-linearized pGCGFP-G418 (a gift from Andrew Pierce, Addgene plasmid #31264) using In-Fusion HD (Takara Bio). ACs C-terminally tagged with muGFP^44^ were generated by removing EGFP from pEGFP-N1 vector by digestion with AgeI and NotI followed by PCR amplification of muGFP (a gift from Mullins lab) and insertion using In-Fusion HD. Next, the generated pmuGFP-N1 vector was linearized with SacI and ApaI and PCR amplified AC3, AC5 and AC9 were inserted using In-Fusion HD. AC5-AC9-Nter-GFP was generated by PCR amplification of AC9 (residues M1-L105) and AC5 (D186-S1261) and insertion into pmuGFP-N1 using In-Fusion HD. AC5-ΔNter-GFP was generated by PCR amplification of AC5 (D192-S1261) and insertion into pmuGFP-N1 using In-Fusion HD. AC9-AC5-Nter-GFP was generated by PCR amplification of AC5 (M1-G185) and AC9 (residues E106-V1353) followed by insertion into pmuGFP-N1 using In-Fusion HD. AC5-AC9-1-35-GFP, AC5-AC9-36-70-GFP and AC5-AC9-71-105-GFP were generated by PCR amplification of AC9 (residues M1-K35, Q36-G70 and G71-L105, respectively) and AC5 (D186-S1261) and insertion into pmuGFP-N1 using In-Fusion HD. To generate AC5-AC9-1-35-LL>AA-GFP, a fragment of AC9 (residues M1-K35) containing the point mutations L9A and L10A (LL>AA) was generated by synthesis (Integrated DNA Technologies, IDT), followed by PCR amplification of AC5 (D192-S1261) and insertion into pmuGFP-N1. AC9-LL>AA-GFP was generated by PCR amplification of AC5-AC9-1-35-LL>AA-GFP (M1-K35) and AC9 (Q36-V1353) and insertion into pmuGFP-N1. pcDNA3.1(+)-ExRai-AKAR2 (a gift from Jin Zhang, Addgene plasmid #161753) was subcloned into pGCGFP-G418. ExRai-AKAR2-NLS and -NES were previously generated in the von Zastrow lab by G. Peng by adding at the C-terminus 2x NLS motifs (SV40, 2x PKKKRKV) or a NES motif (LPPLERLTL), respectively, and subcloned into pGCGFP-G418. AC9-HaloTag was generated by inserting a HaloTag in AC9 extracellular loop 6 between residues E959 and R960. AC9 (M1-E959) was PCR amplified and two DNA fragments containing HaloTag and AC9 R960-V1353 were synthesized (Twist Bioscience), followed by insertion into pGCGFP-G418. PKAcat-GFP was previously generated in the von Zastrow lab by A. Marley by addition of EGFP at the C-terminus and cloning into pCAGGS. FLAG-D1R and PKA RIIβ-mCh were previously generated in the von Zastrow lab by A. Ehrlich and G. Peng, respectively. All constructs were validated by DNA sequencing.

### Immunostaining, confocal and iSIM imaging

Neurons were fixed on DIV 11-12 with 4% paraformaldehyde and 4% sucrose for 10 min at room temperature, and then washed 3 times with PBS. Cells were permeabilized with 0.1% Triton X-100 diluted in PBS for 5 min and washed 3 times with PBS. Cells were blocked in 1% BSA diluted in PBS for 30 min. Primary antibodies were diluted into blocking solution and incubated for 30 min at room temperature. Neurons were washed 3 times with blocking solution and incubated with secondary antibodies diluted in blocking solution. Cells were washed 3 times in blocking solution, labeled with DAPI (Sigma-Aldrich) in PBS for 5 min when indicated, and washed 3 times with PBS. Neurons were imaged by confocal microscopy using a Nikon Ti inverted microscope equipped with a Yokogawa CSU-22 spinning disk unit, a Photometrics Evolve Delta EMCCD camera controlled by NIS-Elements 5.21.03 software and 488, 561, and 640 nm Coherent OBIS lasers. Samples were imaged using an Apo TIRF 100x/1.49 NA oil objective (Nikon), and 0.3 μm z-stacks were acquired. The iSIM was performed with a VT-iSIM super-resolution module on a Nikon Ti-E inverted microscope equipped with a 100x 1.45NA Plan Apo objective and 405, 488, 561, 647 nm lasers. Images were captured using a Hamamatsu Quest camera controlled by μManager 2.1 software (https://www.micro-manager.org), with 0.25 μm z-stacks. Images were deconvoluted using Microvolution plugin^78^ on ImageJ Fiji^79^. Imaris software 10.1 was used to create 3D renderings.

### Cilia localization quantification

To measure the percentage of neurons with ACs in cilia, we imaged ciliated FLAG-D1R-expressing cells and counted the number of cells with visible HA-AC immunoreactivity on the cilium. The cilia enrichment index was measured on unprocessed maximum intensity Z-projection images using ImageJ Fiji^79^. Briefly, regions of interest (ROIs) were manually drawn around the cilium and the transfected cell on the FLAG-D1R or Arl13b channel. Fluorescence intensity of the HA-AC channel was measured inside both ROIs and background subtracted. Cilia enrichment index was determined by dividing the cilia fluorescence by total cell fluorescence.

### Endosome localization quantification

To measure the percentage of neurons with AC-GFP in intracellular puncta, image analysis was performed on unprocessed images on ImageJ Fiji^79^. Maximum intensity Z-projection images of cells transfected with AC-GFP were created. Then, internal puncta were identified by fluorescence values above a threshold set manually and the number of internal particles were quantified using the Analyze Particles command. Cells with 10 or more particles were considered positive and the number of positive neurons was then divided by the total number of cells. To measure endosomal enrichment, image analysis was performed on unprocessed images using custom written scripts in MATLAB (Mathworks, R2020a). Maximum intensity Z-projection images of the AC-GFP channel and the endosomal marker (EEA1/Vps35) channel were created, and a ROI was drawn around the transfected cell on the AC channel. Endosomes within this region were defined by EEA1 or Vps35 fluorescence values above a threshold set manually. AC fluorescence intensity within the created endosomal mask and in the ROI were then measured and background subtracted. Endosomal enrichment index was determined by dividing the endosomal fluorescence by total cell fluorescence.

### Live cell imaging

Neurons were transfected on DIV 8 with Lipofectamine 2000. On the day of imaging, neurons expressing AC9-HaloTag were labeled with 200 nM JF_549_ HaloTag ligand^80^ for 15 min, then washed three times with pre-equilibrated HBS. Neurons were incubated with HBS for 30 min and washed three times. Live cell maging was performed on a Nikon Ti inverted microscope equipped with a Andor Borealis CSU-W1 spinning disk confocal with solid-state 488, 561 and 640 nm lasers (Andor), Plan Fluor VC ×40 1.3 NA and Plan Apo VC ×100 1.4 NA objectives (Nikon) and an Andor Zyla 4.2 sCMOS camera controlled by μManager 2.0 software^81^ (https://www.micro-manager.org). Cells were kept at 37°C in a temperature- and humidity-controlled chamber (Okolab). Dopamine was added after a 1 min baseline, and images were taken every 20 sec for 30 min. Images displayed in **Figure 2c, Extended Data Figure 2a,c, Figure 6c,d** and **Extended Data Figure 4** were processed on ImageJ Fiji^79^ using the Subtract Background and Smooth commands.

### Receptor accumulation into endosomes

Image analysis was performed on unprocessed images using custom written scripts in MATLAB (Mathworks, R2020a). For quantifying receptor accumulation at endosomes, a ROI was drawn around the cell on the receptor channel, refined by thresholding a maximal temporal projection and smoothened. Endosomes within this region were defined by AC9-GFP (**Fig. 2d**) or DsRed-EEA1 (**Extended Data Fig. 2b,d**) fluorescence values above a threshold set manually. Filtering to exclude regions smaller than 3 pixels was applied, remaining objects were smoothened with a dilatation of 1 pixel before an erosion of 1 pixel. The generated mask was applied to the D1R or AC9-GFP channel to measure the fluorescence at the location of marker objects. The average fluorescence in the mask was background subtracted and normalized to baseline (before agonist addition).

### cAMP and PKA activity assay

For the cAMP assay, neurons were transduced with the Green Up cAMP biosensor (Montana Molecular) according to manufacturer’s instructions. On the day of imaging, neurons expressing cADDis/ExRai-AKAR2 biosensors alone or with HA-ACs were washed three times with pre-equilibrated HEPES buffered saline (HBS) solution (NaCl 120mM, KCl 2mM, MgCl_2_ 2mM, CaCl_2_ 2mM, glucose 5mM, HEPES 10mM adjusted to pH 7.4). 10 μM dopamine (Millipore Sigma) was added by bath application after a 1 min baseline, and 10 μM forskolin (Fsk, Millipore Sigma) and 500 μM 3-isobutyl-1-methylxanthine (IBMX, Millipore Sigma) were added 30 min later for 2 min. Images were taken every 20 sec with a 488 nm laser. Image analysis was performed on unprocessed images using ImageJ Fiji^79^. Briefly, ROIs were drawn around individual cells, the average fluorescence in the ROI was background subtracted and normalized to baseline. For cADDIs and non-targeted ExRai-AKAR2, fluorescence was then divided by the average fluorescence of the Fsk/IBMX treatment. Integrated cAMP and PKA responses were measured as the area under the curve of two phases (0-5 min and 5-30 min of dopamine treatment). The peak value corresponds to the maximum fluorescence value between 0-30 min after dopamine addition and normalized to the average HA-AC9 peak value. The plateau phase corresponds to the average fluorescence intensity between 20-30 min of dopamine treatment and normalized to the average HA-AC9 phase value.

### Statistical analysis and reproducibility

All data are shown as mean ± standard error of the mean (SEM). from at least three independent neuronal cultures, and images are representative of at least three independent neuronal cultures. Statistical analyses were performed using Prism 10 (GraphPad) using unpaired *t*-test to determine significance.

## Acknowledgements

We thank L. Lavis, A. Ehrlich, G. Peng and A. Marley for sharing reagents. We thank D. Jullié for assistance with primary neuron culture and for providing MATLAB scripts. We thank C. Dessauer for valuable discussion. We thank R. D. Mullins for the use of the iSIM microscope, S. Lord and A. Charles-Orszag for technical support. We thank members of the von Zastrow laboratory for valuable discussion and feedback on the manuscript. Imaging experiments were primarily carried out in the UCSF Center for Advanced Light Microscopy; we thank K. Herrington and S. Kim for technical support and expertise. CSU-W1 Spinning Disk/High Speed Widefield was supported by S10 Shared Instrumentation grant (1S10OD017993-01A1). We thank the Gladstone Histology and Light Microscopy Core directed by B. Ndjamen for assistance with image analyses. This study was supported by research grants from the National Institutes of Health (grant nos. DA012864, DA010711 and MH120212 to M.v.Z.). L.R. was supported by the European Molecular Biology Organization (ALTF 192-2019).

## Author Contributions

L.R. carried out experiments and analyzed data. L.R. and M.v.Z. conceived the experiments and wrote the manuscript.

## References

1. Zaccolo, M., Zerio, A. & Lobo, M. J. Subcellular Organization of the cAMP Signaling Pathway. Pharmacol Rev 73, 278–309 (2021).

2. Calebiro, D. et al. Persistent cAMP-signals triggered by internalized G-protein-coupled receptors. PLoS Biol 7, e1000172 (2009).

3. Ferrandon, S. et al. Sustained cyclic AMP production by parathyroid hormone receptor endocytosis. Nat Chem Biol 5, 734–742 (2009).

4. Kotowski, S. J., Hopf, F. W., Seif, T., Bonci, A. & von Zastrow, M. Endocytosis promotes rapid dopaminergic signaling. Neuron 71, 278–290 (2011).

5. Feinstein, T. N. et al. Noncanonical control of vasopressin receptor type 2 signaling by retromer and arrestin. J Biol Chem 288, 27849–27860 (2013).

6. Merriam, L. A. et al. Pituitary adenylate cyclase 1 receptor internalization and endosomal signaling mediate the pituitary adenylate cyclase activating polypeptide-induced increase in guinea pig cardiac neuron excitability. J Neurosci 33, 4614–4622 (2013).

7. Tsvetanova, N. G. & von Zastrow, M. Spatial encoding of cyclic AMP signaling specificity by GPCR endocytosis. Nat Chem Biol 10, 1061–1065 (2014).

8. Lyga, S. et al. Persistent cAMP Signaling by Internalized LH Receptors in Ovarian Follicles. Endocrinology 157, 1613–1621 (2016).

9. Godbole, A., Lyga, S., Lohse, M. J. & Calebiro, D. Internalized TSH receptors en route to the TGN induce local Gs-protein signaling and gene transcription. Nat Commun 8, 443 (2017).

10. Jensen, D. D. et al. Neurokinin 1 receptor signaling in endosomes mediates sustained nociception and is a viable therapeutic target for prolonged pain relief. Sci Transl Med 9, eaal3447 (2017).

11. Jimenez-Vargas, N. N. et al. Endosomal signaling of delta opioid receptors is an endogenous mechanism and therapeutic target for relief from inflammatory pain. Proc Natl Acad Sci U S A 117, 15281–15292 (2020).

12. White, A. D. et al. Spatial bias in cAMP generation determines biological responses to PTH type 1 receptor activation. Sci Signal 14, eabc5944 (2021).

13. Zaccolo, M. Phosphodiesterases and compartmentalized cAMP signalling in the heart. Eur J Cell Biol 85, 693–697 (2006).

14. Kritzer, M. D., Li, J., Dodge-Kafka, K. & Kapiloff, M. S. AKAPs: the architectural underpinnings of local cAMP signaling. J Mol Cell Cardiol 52, 351–358 (2012).

15. Conti, M., Mika, D. & Richter, W. Cyclic AMP compartments and signaling specificity: role of cyclic nucleotide phosphodiesterases. J Gen Physiol 143, 29–38 (2014).

16. Wild, A. R. & Dell’Acqua, M. L. Potential for therapeutic targeting of AKAP signaling complexes in nervous system disorders. Pharmacol Ther 185, 99–121 (2018).

17. Dessauer, C. W. et al. International Union of Basic and Clinical Pharmacology. CI. Structures and Small Molecule Modulators of Mammalian Adenylyl Cyclases. Pharmacol Rev 69, 93–139 (2017).

18. Cooper, D. M. F. & Crossthwaite, A. J. Higher-order organization and regulation of adenylyl cyclases. Trends Pharmacol Sci 27, 426–431 (2006).

19. Johnstone, T. B., Agarwal, S. R., Harvey, R. D. & Ostrom, R. S. cAMP Signaling Compartmentation: Adenylyl Cyclases as Anchors of Dynamic Signaling Complexes. Mol Pharmacol 93, 270–276 (2018).

20. Dessauer, C. W. Adenylyl cyclase--A-kinase anchoring protein complexes: the next dimension in cAMP signaling. Mol Pharmacol 76, 935–941 (2009).

21. Baldwin, T. A. & Dessauer, C. W. Function of Adenylyl Cyclase in Heart: the AKAP Connection. J Cardiovasc Dev Dis 5, 2 (2018).

22. Cheng, H. & Farquhar, M. G. Presence of adenylate cyclase activity in Golgi and other fractions from rat liver. I. Biochemical determination. J Cell Biol 70, 660–670 (1976).

23. Cheng, H. & Farquhar, M. G. Presence of adenylate cyclase activity in Golgi and other fractions from rat liver. II. Cytochemical localization within Golgi and ER membranes. J Cell Biol 70, 671–684 (1976).

24. Cancino, J. et al. Control systems of membrane transport at the interface between the endoplasmic reticulum and the Golgi. Dev Cell 30, 280–294 (2014).

25. Jean-Alphonse, F. G. et al. β2-adrenergic receptor control of endosomal PTH receptor signaling via Gβγ. Nat Chem Biol 13, 259–261 (2017).

26. Lazar, A. M. et al. G protein-regulated endocytic trafficking of adenylyl cyclase type 9. Elife 9, e58039 (2020).

27. Boczek, T. et al. Regulation of Neuronal Survival and Axon Growth by a Perinuclear cAMP Compartment. J Neurosci 39, 5466–5480 (2019).

28. Shelly, M. et al. Local and long-range reciprocal regulation of cAMP and cGMP in axon/dendrite formation. Science 327, 547–552 (2010).

29. Gorshkov, K. et al. AKAP-mediated feedback control of cAMP gradients in developing hippocampal neurons. Nat Chem Biol 13, 425–431 (2017).

30. Averaimo, S. et al. A plasma membrane microdomain compartmentalizes ephrin-generated cAMP signals to prune developing retinal axon arbors. Nat Commun 7, 12896 (2016).

31. Maiellaro, I., Lohse, M. J., Kittel, R. J. & Calebiro, D. cAMP Signals in Drosophila Motor Neurons Are Confined to Single Synaptic Boutons. Cell Rep 17, 1238–1246 (2016).

32. Calebiro, D. & Maiellaro, I. cAMP signaling microdomains and their observation by optical methods. Front Cell Neurosci 8, 350 (2014).

33. Oliveira, R. F., Kim, M. & Blackwell, K. T. Subcellular location of PKA controls striatal plasticity: stochastic simulations in spiny dendrites. PLoS Comput Biol 8, e1002383 (2012).

34. Brandon, E. P. et al. Defective motor behavior and neural gene expression in RIIbeta-protein kinase A mutant mice. J Neurosci 18, 3639–3649 (1998).

35. Matamales, M. & Girault, J.-A. Signaling from the cytoplasm to the nucleus in striatal medium-sized spiny neurons. Front Neuroanat 5, 37 (2011).

36. Matsuoka, I., Suzuki, Y., Defer, N., Nakanishi, H. & Hanoune, J. Differential expression of type I, II, and V adenylyl cyclase gene in the postnatal developing rat brain. J Neurochem 68, 498–506 (1997).

37. Sanabra, C. & Mengod, G. Neuroanatomical distribution and neurochemical characterization of cells expressing adenylyl cyclase isoforms in mouse and rat brain. J Chem Neuroanat 41, 43–54 (2011).

38. Lee, K.-W. et al. Impaired D2 dopamine receptor function in mice lacking type 5 adenylyl cyclase. J Neurosci 22, 7931–7940 (2002).

39. Iwamoto, T. et al. Disruption of type 5 adenylyl cyclase negates the developmental increase in Galphaolf expression in the striatum. FEBS Lett 564, 153–156 (2004).

40. Bishop, G. A., Berbari, N. F., Lewis, J. & Mykytyn, K. Type III adenylyl cyclase localizes to primary cilia throughout the adult mouse brain. J Comp Neurol 505, 562–571 (2007).

41. Kheirbek, M. A., Beeler, J. A., Ishikawa, Y. & Zhuang, X. A cAMP pathway underlying reward prediction in associative learning. J Neurosci 28, 11401–11408 (2008).

42. Kheirbek, M. A., Beeler, J. A., Chi, W., Ishikawa, Y. & Zhuang, X. A molecular dissociation between cued and contextual appetitive learning. Learn Mem 17, 148–154 (2010).

43. Berbari, N. F., Bishop, G. A., Askwith, C. C., Lewis, J. S. & Mykytyn, K. Hippocampal neurons possess primary cilia in culture. J Neurosci Res 85, 1095–1100 (2007).

44. Scott, D. J. et al. A Novel Ultra-Stable, Monomeric Green Fluorescent Protein For Direct Volumetric Imaging of Whole Organs Using CLARITY. Sci Rep 8, 667 (2018).

45. Hilger, D., Masureel, M. & Kobilka, B. K. Structure and dynamics of GPCR signaling complexes. Nat Struct Mol Biol 25, 4–12 (2018).

46. Puri, N. M., Romano, G. R., Lin, T.-Y., Mai, Q. N. & Irannejad, R. The organic cation transporter 2 regulates dopamine D1 receptor signaling at the Golgi apparatus. Elife 11, e75468 (2022).

47. Stubbs, T. et al. Disruption of dopamine receptor 1 localization to primary cilia impairs signaling in striatal neurons. J Neurosci 42, 6692–6705 (2022).

48. Marley, A. & von Zastrow, M. DISC1 regulates primary cilia that display specific dopamine receptors. PLoS One 5, e10902 (2010).

49. Domire, J. S. et al. Dopamine receptor 1 localizes to neuronal cilia in a dynamic process that requires the Bardet-Biedl syndrome proteins. Cell Mol Life Sci 68, 2951–2960 (2011).

50. Vickery, R. G. & von Zastrow, M. Distinct dynamin-dependent and -independent mechanisms target structurally homologous dopamine receptors to different endocytic membranes. J Cell Biol 144, 31–43 (1999).

51. Dumartin, B., Caillé, I., Gonon, F. & Bloch, B. Internalization of D1 dopamine receptor in striatal neurons in vivo as evidence of activation by dopamine agonists. J Neurosci 18, 1650–1661 (1998).

52. Traub, L. M. & Bonifacino, J. S. Cargo recognition in clathrin-mediated endocytosis. Cold Spring Harb Perspect Biol 5, a016790 (2013).

53. Zhang, J.-F. et al. An ultrasensitive biosensor for high-resolution kinase activity imaging in awake mice. Nat Chem Biol 17, 39–46 (2021).

54. Cadd, G. & McKnight, G. S. Distinct patterns of cAMP-dependent protein kinase gene expression in mouse brain. Neuron 3, 71–79 (1989).

55. Ilouz, R. et al. Isoform-specific subcellular localization and function of protein kinase A identified by mosaic imaging of mouse brain. Elife 6, e17681 (2017).

56. Ventra, C. et al. The differential response of protein kinase A to cyclic AMP in discrete brain areas correlates with the abundance of regulatory subunit II. J Neurochem 66, 1752–1761 (1996).

57. Irannejad, R. et al. Functional selectivity of GPCR-directed drug action through location bias. Nat Chem Biol 13, 799–806 (2017).

58. Nigg, E. A., Schäfer, G., Hilz, H. & Eppenberger, H. M. Cyclic-AMP-dependent protein kinase type II is associated with the Golgi complex and with centrosomes. Cell 41, 1039–1051 (1985).

59. Peng, G. E., Pessino, V., Huang, B. & von Zastrow, M. Spatial decoding of endosomal cAMP signals by a metastable cytoplasmic PKA network. Nat Chem Biol 17, 558–566 (2021).

60. Nigg, E. A., Hilz, H., Eppenberger, H. M. & Dutly, F. Rapid and reversible translocation of the catalytic subunit of cAMP-dependent protein kinase type II from the Golgi complex to the nucleus. EMBO J 4, 2801–2806 (1985).

61. Mavillard, F., Hidalgo, J., Megias, D., Levitsky, K. L. & Velasco, A. PKA-mediated Golgi remodeling during cAMP signal transmission. Traffic 11, 90–109 (2010).

62. Willette, B. K. A., Zhang, J.-F., Zhang, J. & Tsvetanova, N. G. Endosome positioning coordinates spatially selective GPCR signaling. Nat Chem Biol (2023) doi:10.1038/s41589-023-01390-7.

63. Hagiwara, M. et al. Coupling of hormonal stimulation and transcription via the cyclic AMP-responsive factor CREB is rate limited by nuclear entry of protein kinase A. Mol Cell Biol 13, 4852–4859 (1993).

64. Kandel, E. R. The molecular biology of memory: cAMP, PKA, CRE, CREB-1, CREB-2, and CPEB. Mol Brain 5, 14 (2012).

65. Altarejos, J. Y. & Montminy, M. CREB and the CRTC co-activators: sensors for hormonal and metabolic signals. Nat Rev Mol Cell Biol 12, 141–151 (2011).

66. Zhang, H., Kong, Q., Wang, J., Jiang, Y. & Hua, H. Complex roles of cAMP-PKA-CREB signaling in cancer. Exp Hematol Oncol 9, 32 (2020).

67. Schmidt, U., Pilgrim, C. & Beyer, C. Differentiative effects of dopamine on striatal neurons involve stimulation of the cAMP/PKA pathway. Mol Cell Neurosci 11, 9–18 (1998).

68. Fienberg, A. A. & Greengard, P. The DARPP-32 knockout mouse. Brain Res Brain Res Rev 31, 313–319 (2000).

69. Fienberg, A. A. et al. DARPP-32: regulator of the efficacy of dopaminergic neurotransmission. Science 281, 838–842 (1998).

70. Choi, E. J., Xia, Z. & Storm, D. R. Stimulation of the type III olfactory adenylyl cyclase by calcium and calmodulin. Biochemistry 31, 6492–6498 (1992).

71. Taussig, R., Tang, W. J., Hepler, J. R. & Gilman, A. G. Distinct patterns of bidirectional regulation of mammalian adenylyl cyclases. J Biol Chem 269, 6093–6100 (1994).

72. Baldwin, T. A., Li, Y., Brand, C. S., Watts, V. J. & Dessauer, C. W. Insights into the Regulatory Properties of Human Adenylyl Cyclase Type 9. Mol Pharmacol 95, 349–360 (2019).

73. O’Banion, C. P., Vickerman, B. M., Haar, L. & Lawrence, D. S. Compartmentalized cAMP Generation by Engineered Photoactivated Adenylyl Cyclases. Cell Chem Biol 26, 1393-1406.e7 (2019).

74. Ferrell, J. E. What do scaffold proteins really do? Sci STKE 2000, pe1 (2000).

75. Prinz, W. A., Toulmay, A. & Balla, T. The functional universe of membrane contact sites. Nat Rev Mol Cell Biol 21, 7–24 (2020).

76. Antoni, F. A. et al. Ca2+/calcineurin-inhibited adenylyl cyclase, highly abundant in forebrain regions, is important for learning and memory. J Neurosci 18, 9650–9661 (1998).

77. Ventimiglia, R. & Lindsay, R. M. Rat Striatal Neurons in Low-Density, Serum-Free Culture. in Culturing Nerve Cells (eds. Banker, G. & Goslin, K.) 371–394 (The MIT Press, 1998). doi:10.7551/mitpress/4913.003.0021.

78. Bruce, M. A. & Butte, M. J. Real-time GPU-based 3D Deconvolution. Opt Express 21, 4766–4773 (2013).

79. Schindelin, J. et al. Fiji: an open-source platform for biological-image analysis. Nat Methods 9, 676–682 (2012).

80. Grimm, J. B., Brown, T. A., English, B. P., Lionnet, T. & Lavis, L. D. Synthesis of Janelia Fluor HaloTag and SNAP-Tag Ligands and Their Use in Cellular Imaging Experiments. Methods Mol Biol 1663, 179–188 (2017).

81. Edelstein, A. D. et al. Advanced methods of microscope control using μManager software. J Biol Methods 1, e10 (2014).

