## Supplemental figures for "Spatial organization of adenylyl cyclase and its impact on dopamine signaling in neurons"

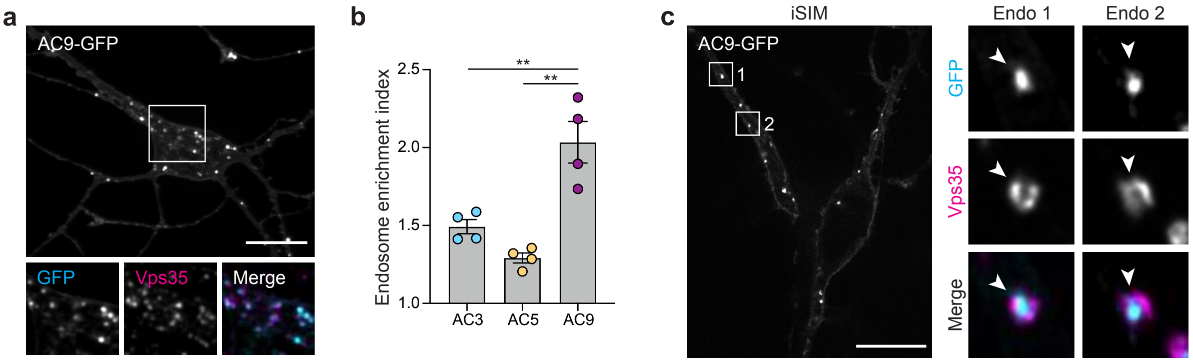


**Extended Data Fig 1. AC9 localizes to endosomes. a,** Maximum intensity Z-projection of confocal microscopy images of MSNs expressing AC9-GFP and stained for endosomal marker Vps35. **b,** Endosome enrichment index calculated as a ratio of AC-GFP fluorescence intensity at EEA1 positive endosomes divided by total cell fluorescence. Data are shown as mean ± s.e.m from $n$ = 4. **c,** Representative iSIM images of MSN expressing AC9-GFP and stained for Vps35. Arrowheads indicate the endosomal membrane. Data represent biological replicates and are shown as individual data points or mean ± s.e.m. Scale bars are 10 μm. **$P$<0.01 by unpaired two-tailed Student's $t$-test.

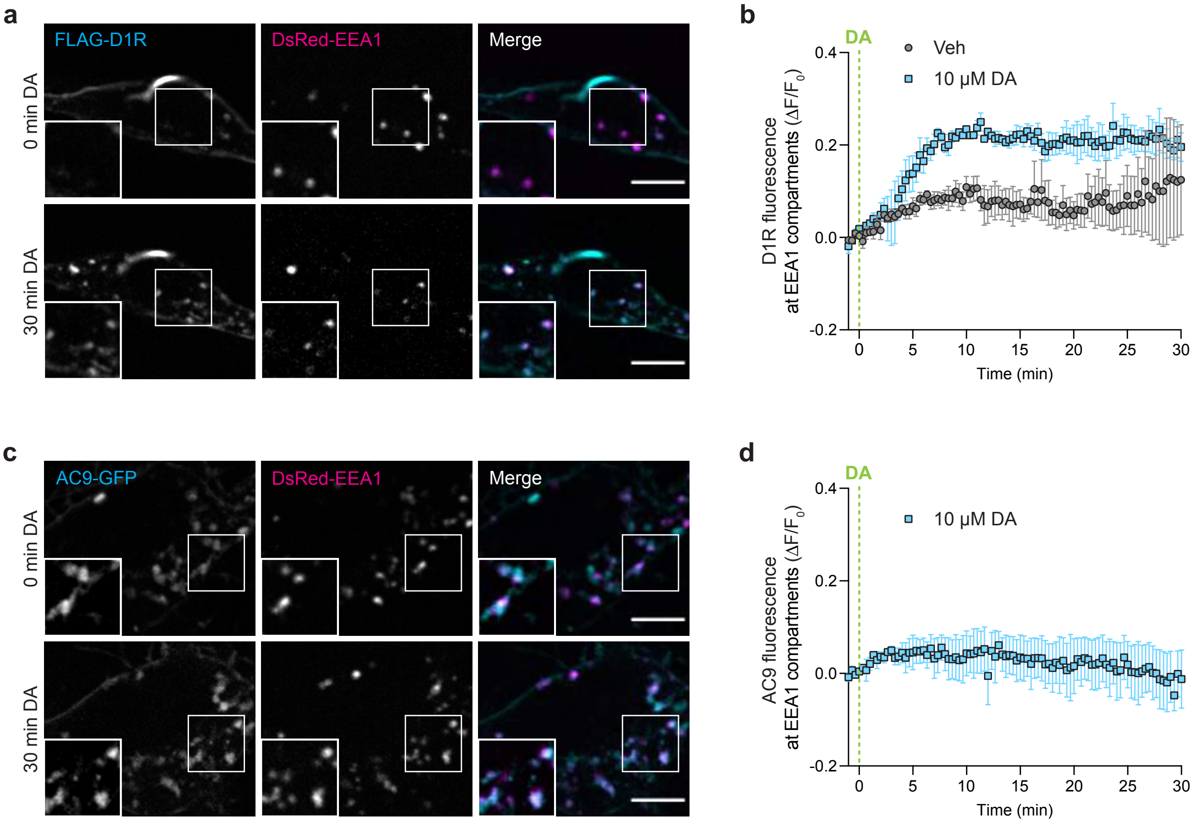


**Extended Data Fig 2. Dopamine 1 receptor and AC9 localize to EEA1 positive endosomes. a,** Representative live cell spinning disk confocal images of MSNs transfected with FLAG-D1R and DsRed-EEA1 and treated with 10 μM dopamine (DA) at 0 min. Surface FLAG-D1R was labeled with Alexa Fluor 555-coupled anti-FLAG antibody for 15 min before imaging. **b,** Quantification of surface labeled FLAG-D1R accumulation at segmented EEA1-positive endosomes after vehicle (Veh) or 10 μM DA addition. Data are shown as mean ± s.e.m from $n$ = 3. **c,** Representative live cell spinning disk confocal images of MSNs transfected with AC9-GFP and DsRed-EEA1 and treated with 10 μM DA at 0 min. **d,** Quantification of AC9-GFP accumulation at segmented EEA1-positive endosomes after 10 μM DA addition. Data are shown as mean ± s.e.m from $n$ = 3. Scale bars are 5 μm.

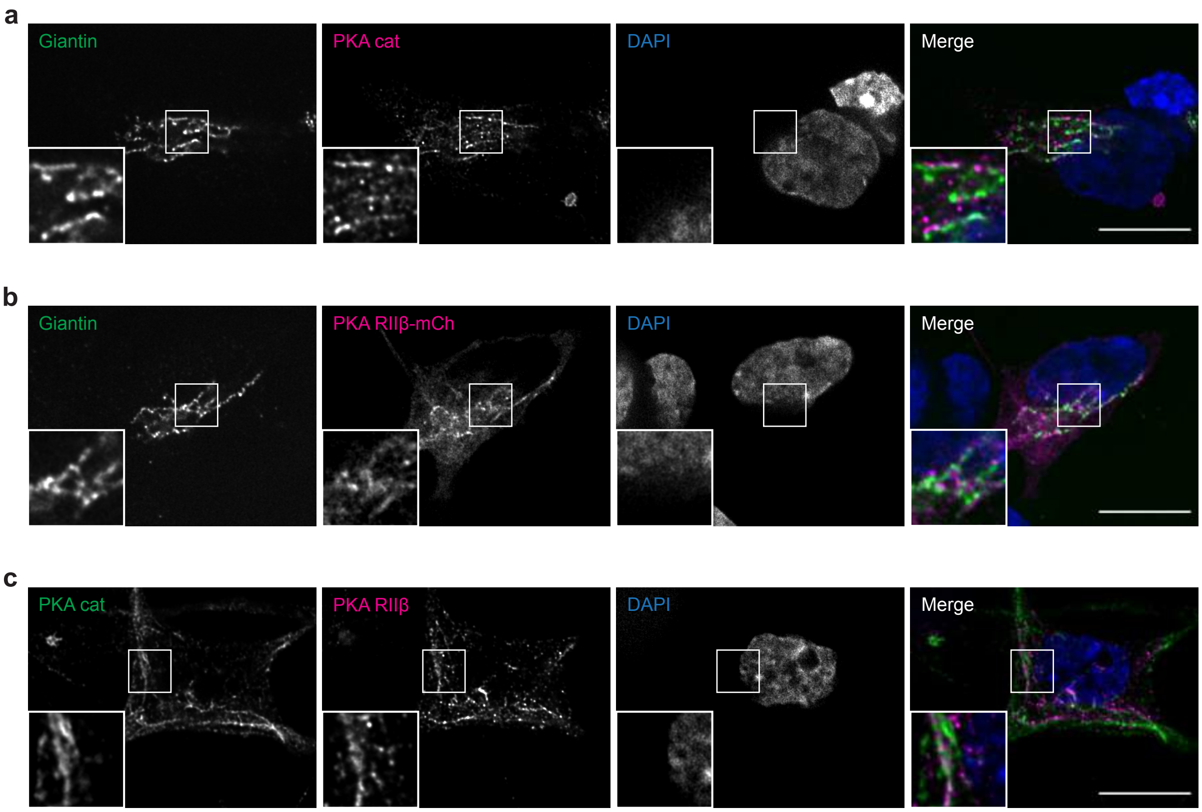


**Extended Data Fig 3. PKA subunits localization at the Golgi. a,** Representative iSIM images of MSN stained for the Golgi marker giantin and endogenous PKA cat. **b,** Representative iSIM images of MSNs expressing PKA RIIβ-mCh and stained for giantin. **c,** Representative iSIM images of MSN stained for endogenous PKA cat and PKA RIIβ. Nuclei were stained with DAPI. Scale bars are 10 μm.

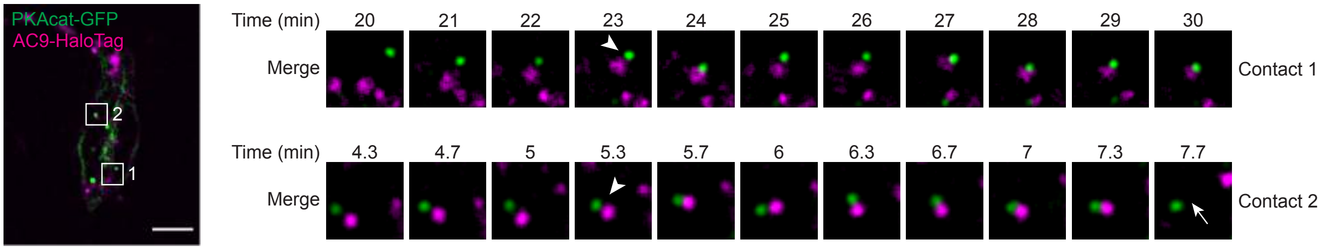


**Extended Data Fig 4. Dynamic contacts between PKA cat puncta and AC9 containing endosomes.** Spinning-disk confocal images from a time series of neurons expressing PKAcat-GFP and AC9-HaloTag and treated with 10 μM DA at t = 0 min. PKA cat puncta and AC9-containing endosomes form close contacts (arrowheads) for several minutes then separate (arrow). Scale bar is 5 μm.

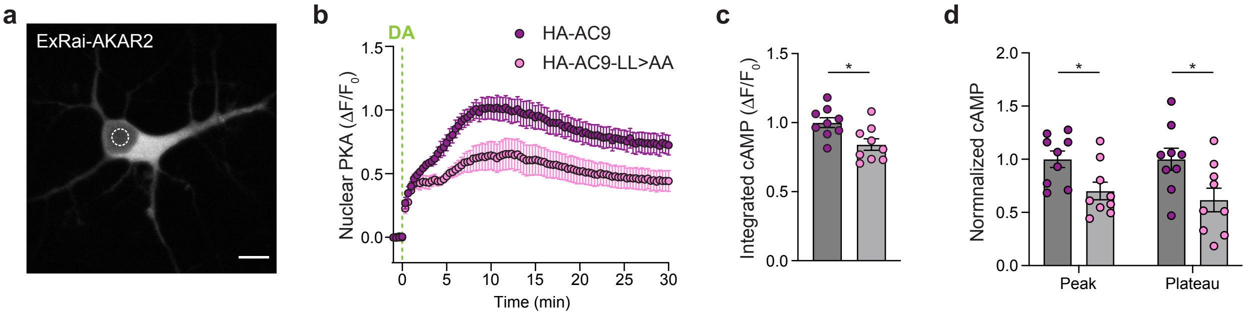


**Extended Data Fig 5. Mutating AC9 dileucine motif impacts PKA activity in the nucleus. a,** Spinning-disk confocal representative image of MSN expressing ExRai-AKAR2. A ROI is drawn inside the nucleus (white dotted circle) to measure PKA activity within the nucleus over time. Scale bar = 10 µm. **b,** Kinetics of nuclear PKA activity over time in MSNs coexpressing ExRai-AKAR2 and HA-AC9 (purple, *n* = 9) or HA-AC9-LL>AA (pink, *n* = 9) and treated with 10 µM dopamine (DA) The ΔF/F0 was measured every 20 sec. **c,** Integrated nuclear PKA signal was calculated as the area under the curve and normalized to the average HA-AC9 value. **d,** Peak and plateau values were calculated as the maximum ΔF/F0 (peak) and the average of 20-30 min values (plateau) and normalized to the HA-AC9 value. Data represent biological replicates and are shown as mean ± s.e.m. **P*<0.05 by unpaired two-tailed Student's *t*-test.

**Supplementary Video 1**
Movie of the 3D rendering of MSN expressing AC9-GFP (green) stained for endogenous PKA cat (magenta) and stained with DAPI (blue) to label the nucleus. The video shows extensive regions of close apposition between AC9-containing endosomes and Golgi-associated PKA cat stores close to the nucleus.

**Supplementary Video 1**
Movie of the 3D rendering of MSN expressing AC9-GFP (green) and stained for endogenous PKA RIIβ (magenta). Nucleus was stained with DAPI (blue). The video shows close proximity or contact between AC9-positive endosomes and Golgi-associated PKA RIIβ compartments next to the nucleus.

**Supplementary Video 3**
Movie of a live confocal image series of MSN expressing PKAcat-GFP (green) and AC9-HaloTag (magenta) and treated with 10 μM dopamine (DA). The video shows PKA cat puncta and AC9-containing endosomes moving together for several minutes, indicating close contact. Scale bar, 1 μm.
